# Membrane pore energetics and the pathways to membrane rupture

**DOI:** 10.1101/2020.06.29.178988

**Authors:** Dong An, Sathish Thiyagarajan, Egor Antipov, Brett Alcott, Ben O’Shaughnessy

## Abstract

Biological membranes owe their strength and low permeability to the phospholipid bilayers at their core. Membrane strength is determined by the energetics and dynamics of membrane pores, whose tension-dependent nucleation and growth leads to rupture. Creation of nanoscale membrane pores is central to exocytosis, trafficking and other processes fundamental to life that require breaching of secure plasma or organelle membranes, and is the basis for biotechnologies using drug delivery, delivery of genetic material for gene editing and antimicrobial peptides. A prevailing view from seminal electroporation and membrane rupture studies is that pore growth and bilayer rupture are controlled by macroscopically long-lived metastable defect states that precede fully developed pores. It was argued that defect nucleation becomes rate-limiting at high tensions, explaining the exponential tension-dependence of rupture times [E. Evans et al., *Biophys. J.* 85, 2342-2350 (2003)]. Here we measured membrane pore free energies and bilayer rupture using highly coarse-grained simulations that probe very long time scales. We find no evidence of metastable pore states. At lower tensions, small hydrophobic pores mature into large hydrophilic pores on the pathway to rupture, with classical tension dependence of rupture times. Above a critical tension membranes rupture directly from a small hydrophobic pore, and rupture times depend exponentially on tension. Thus, we recover the experimentally reported regimes, but the origin of the high tension exponential regime is unrelated to macroscopically long-lived pre-pore defects. It arises because hydrophilic pores cannot exist above a critical tension, leading to radically altered pore dynamics and rupture kinetics.

## Introduction

Compartmentalization was a vital development for the emergence of life, made possible by the unique material properties of phospholipid bilayers (1). Three broad properties make them ideal boundaries for cells and subcellular compartments: softness, impermeability and strength. Softness together with impermeability allows facile shape change while restricting the passage of macromolecules, polar or charged small molecules and ions, so that specific biochemical conditions can be maintained in specialized compartments (2, 3). Phospholipid bilayers also endow biological membranes with considerable strength, needed to maintain the substantial osmotic pressures that accompany differential chemical conditions and for safe installation of proteins and other macromolecules for selective transport, signalling and other functions (4).

Many processes fundamental to life require these secure membrane boundaries to be breached by creation of pores, or even irreversibly lysed. For trafficking, or for secretion of neurotransmitters or hormones by exocytosis, cells connect compartments via fusion pores by puncturing and fusing the membranes (5). The pathways to membrane fusion remain controversial, but in some cases fusion is completed by lysis of a hemifusion diaphragm intermediate, likely through formation and growth of a simple pore (2, 6–9). Similar challenges are faced by pathogens such as influenza, HIV and SARS coronaviruses, which introduce their genomes through fusion pores following cell entry by fusion of the viral and host membranes (10, 11).

Formation of nanoscale membrane pores is the mechanism of reversible electrical breakdown of membranes (12), and is exploited by a number of biotechnologies and antimicrobial strategies (13–16). Delivery of drugs or genetic material can be achieved by opening simple pores using electroporation (14, 17), detergents (18), or cell-penetrating peptides (19, 20). For example, the cell-penetrating peptide nona-arginine (R9) facilitates Cas9 entry in the CRISPR-Cas9 gene editing system (15, 16). Naturally produced antimicrobial peptides are used by multicellular organisms to make pores in microbial membranes (21) and have potential as therapeutic agents (22, 23).

Simple pore energetics and dynamics are fundamentally related to bilayer strength. In the classical model the pore of radius *R* has free energy *F* = 2*πτR* − *γπR*^2^, quantifying how membrane tension *γ* promotes rupture by pore formation and expansion while pore line tension *τ* opposes growth (24). In this picture, a pore fluctuation large enough to surmount the barrier *F*_max_ = *πτ*^2^/*γ* will grow the pore without bound and rupture the membrane.

Evans et al. measured membrane rupture times by ramping up giant unilamellar vesicle (GUV) tensions at variable rates (25). A pore kinetics model was developed, featuring a nanoscale (~ 0.6 nm radius) metastable defect state that preceded the classical membrane pore. However a very long ~ 1s defect state lifetime was required in order for the model to fit their high tension data which implied exponential dependence of lysis times on tension (25). An earlier study of black lipid membrane (BLM) electroporation similarly suggested that pore opening kinetics involved metastable pre-pore defect states with extremely long lifetimes 0.1-1 s (26). The possibility of metastable pre-pore states was suggested by other theoretical models and atomistic simulations (12, 27, 28).

Here we studied simple pores and bilayer rupture in simulated phospholipid bilayers with highly coarse grained lipids. This approach allowed us to probe timescales up to tens of milliseconds, and pore sizes up to the limit of stability. Using an umbrella sampling method to measure pore free energies, we find no evidence of metastable pore states at any tension. At low tensions membrane rupture proceeds by growth of a hydrophobic pore into a hydrophilic pore, with classical dependence of lysis times on tension. Above a critical tension membranes rupture directly from a small hydrophobic pore, and lysis times depend on tension in a non-classical manner. Thus, we recover the two regimes reported by Evans et al. (25), but we find the origin of the non-classical high tension regime is unrelated to macroscopically long-lived pre-pore defects. Rather, it arises because stable hydrophilic pores cannot exist above a critical tension, leading to radically altered pore dynamics and rupture kinetics.

## Results

### Model

#### Coarse-grained simulations of phospholipid bilayers

Each lipid is a linear sequence with one hydrophilic head bead (h), and three hydrophobic tail beads (t), interacting according to the force field developed in refs. (29–31). The potential between two beads separated by *r* is *V*_x_ = *V*_rep_(*r*/b_x_) + *α*_x_*V*_attr_(*r*/b_x_) where x is hh, ht or tt. *V*_rep_ is the repulsive Weeks-Chandler-Andersen potential, *V*_attr_ is an attractive potential, *b*_x_ sets the interaction length scale, and *α*_hh_ = *α*_ht_ = 0, *α*_tt_ = 1 (see Supplementary Information). Thus when close enough all beads repel, while tail beads also attract one another. We choose *b*_tt_ = *σ* and *b*_hh_ = *b*_ht_ = 0.95 *σ*, where *σ* and 0.95*σ* are approximately the diameter of a tail and head bead, respectively. To reproduce the experimentally observed bilayer thickness of 5 nm for pure phosphatidylcholine (32), we chose *σ* = 0.88 nm. We simulated 88 × 88 nm bilayers, except for tension-density relationship (Fig. S1) and membrane rupture simulations which used 23 × 23 nm bilayers.

Simulations were run using the HOOMD-blue simulation toolkit in the NVT ensemble (33, 34). Lipids were initially placed in a uniformly spaced grid at the density corresponding to the desired tension from our tension-density relation from separate simulations (Fig. S1). Data were collected following an equilibration period lasting 20 μs (except for tension of 2.5 pN nm^−1^, where equilibration lasted 10 μs) and the tension was taken as the time-averaged value in that simulation. To translate one simulation timestep into physical time Δ*t* we equated the previously measured lipid diffusivity *D* = 3 × 10^−4^ *σ*^2^/ Δ*t* (29) to a typical experimental value, *D* = 1 μm^2^ s^−1^(35), giving *Δt* = 0.2 ns.

#### Pore size constraint

To study a pore of a given radius R we used an umbrella sampling approach, with a simple bias potential constraining pore size. A cylinder oriented perpendicular to the bilayer has hard core radius *R*_∞_ and a halo of thickness *ϵ* = 0.1 nm (Fig. 1), so that any bead whose center of gravity lies a distance r from the cylinder axis is repelled according to the potential

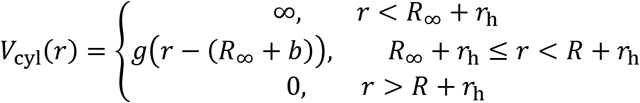

where *g*(*x*) = 0.6*k*_B_*T* [(*ϵ/x*)^12^ − 2(*ϵ/x*)^6^ + 1] is the halo of width *ϵ*. Here *r*_h_ = 2^1/6^*b*_hh_/2 is the head bead radius (see Supplementary Information). To equilibrate a pore the bilayer is equilibrated for 20 μs in the presence of the cylinder. As the halo potential increases rapidly, the cylinder has effective radius *R* = *R*_∞_ + *ϵ*, and provides a constraint that the pore edge can never intersect a cylindrical region of radius *R*.

**Figure 1.**
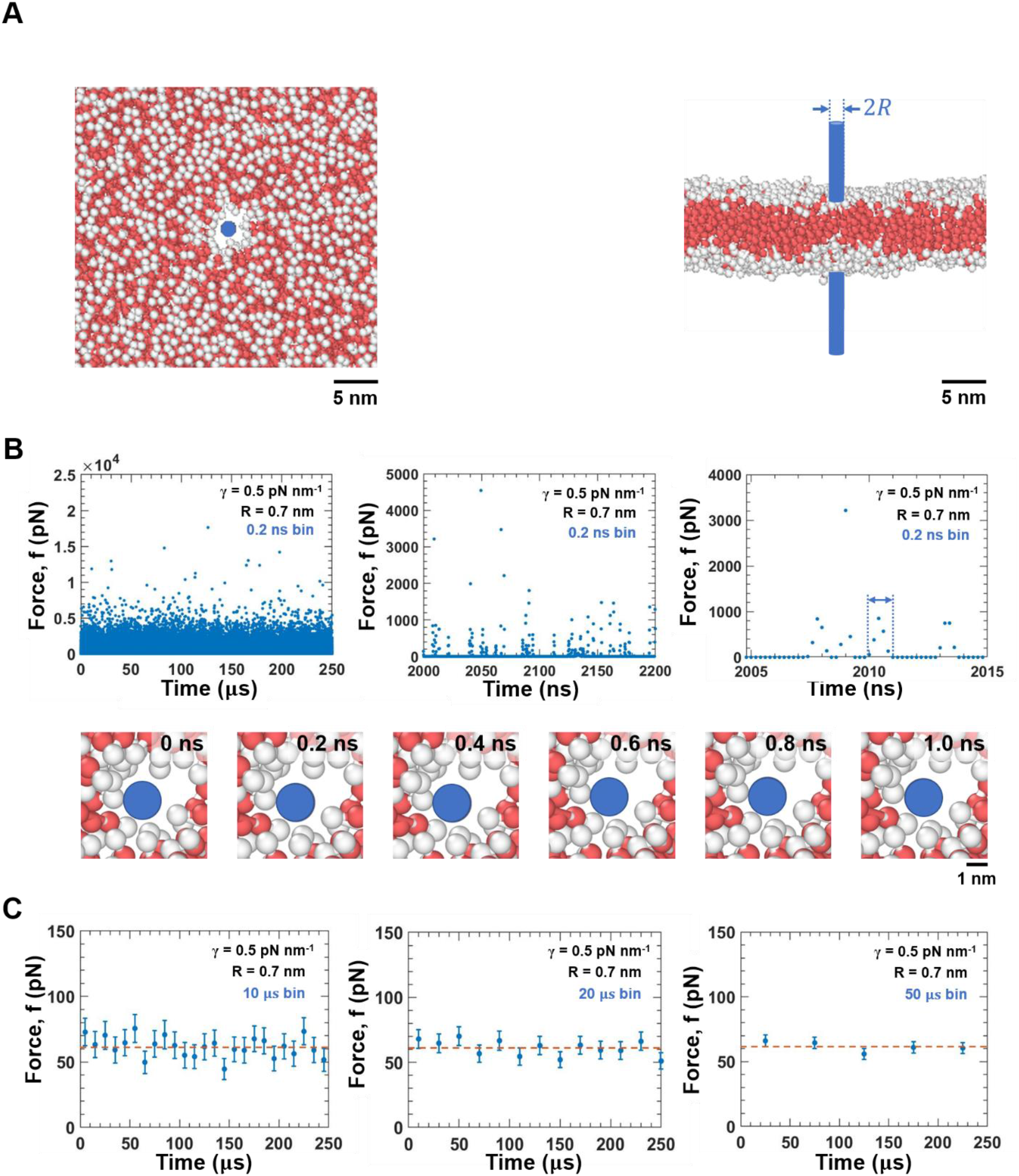
A phospholipid bilayer pore exerts a highly intermittent force of tens of pN. **(A)** Top and side views of a typical simulated 88 × 88 nm bilayer at tension *γ* = 1 pN nm^−1^ containing a pore imposed by a cylinder of radius *R* = 0.8 nm. Lipids are represented by the coarse grained implicit solvent force field of refs. (29–31), with one hydrophilic head bead (white) and three hydrophobic tail beads (red) per lipid. **(B)** Force vs time exerted by a pore on a cylinder of radius 0.7 nm in a bilayer at tension 0.5 pN nm^−1^. The force is plotted for every simulation timestep of duration 0.2 ns, at three time resolutions. The highly intermittent force shows bursts separated by ~ 20 ns silent periods. Bottom: snapshots of the pore during a typical force burst arrowed in the top right panel. Times indicated relative to the start of the burst. **(C)** Binned force vs time of **(B)**, with 10, 20, and 50 μs bins. Time averaged force is 60.7 pN (dashed red line). Bars: S.E.M.

#### Measuring the force on the cylinder

We define the instantaneous net radial force *f*acting on the cylinder to be the sum of the inward radial components of forces exerted by all beads on the cylinder at that timestep. In simulations we measure the force every timestep of duration 0.2 ns and then time average. We found the force is highly intermittent, with force bursts separated by silent periods lasting ~ 20 ns (Fig. 1). Thus, if the force was zero for 200 ns, we deemed this to signal that the pore had irreversibly expanded and the bilayer lysed. Subsequent force measurements were excluded from further analysis. For each pore radius and membrane tension we performed 30 simulations lasting ~ 5-50 μs.

### A membrane pore exerts a highly intermittent force

Our principal goal is to measure the simple pore free energy *F*(*R*) as a function of pore radius *R* for a range of membrane tensions. We simulated bilayers with simple pores imposed by the presence of a cylinder of radius *R* that repels head and tail lipid beads, Fig. 1A. We first calculated the sum of the radial components of the force exerted by the pore on the cylinder (the ‘net radial force’ on the cylinder), whose time average *f*(*R*) will be integrated to yield the free energy (see below).

The instantaneous force was stochastic and highly intermittent. A typical force history for a cylinder of radius 0.7 nm in a bilayer at tension 0.5 pN nm^−1^ is shown in Fig. 1B. As the pore fluctuated in shape, pore collisions with the cylinder produced force bursts of ~ 1ns, separated by silent periods of ~ 10 ns with zero force during which the pore shape changed very little. Substantial shape changes occurred over ~ 1 − 10 μs, the pore configuration memory time. With increasing bin size (10 to 20 to 50 μs), the plotted force converged to the time average of ~ 60 pN with decreasing fluctuations. For other cylinder radii and membrane tensions, the forces exerted by pores showed the same qualitative features (Fig. S2).

### Small hydrophobic pores exert high force, large hydrophilic pores exert low force

The force exerted by a membrane pore depended strongly on its size, and was correlated with its type: small pores were hydrophobic and exerted large forces, while large pores had hydrophilic character and exerted much lower forces (Fig. 2). At a given membrane tension, the time-averaged force versus pore radius was large at small pore sizes, then dropped sharply, followed by a gradual decrease with increasing pore size. For example, at membrane tension *γ* = 1 pN nm^−1^ the spike corresponded to pores with radii up to ~ 0.4 nm, with a peak force ~ 230 pN at radius ~ 0.15 nm, Fig. 2A (forces are mean values over ~ 20-50 μs simulation times). The force dropped to ~ 60 pN for larger pores, then slowly decreasing with a slope close to the value −2πγ = −6.3 pN nm^−1^ predicted by the classical theory, Fig. 2A.

**Figure 2.**
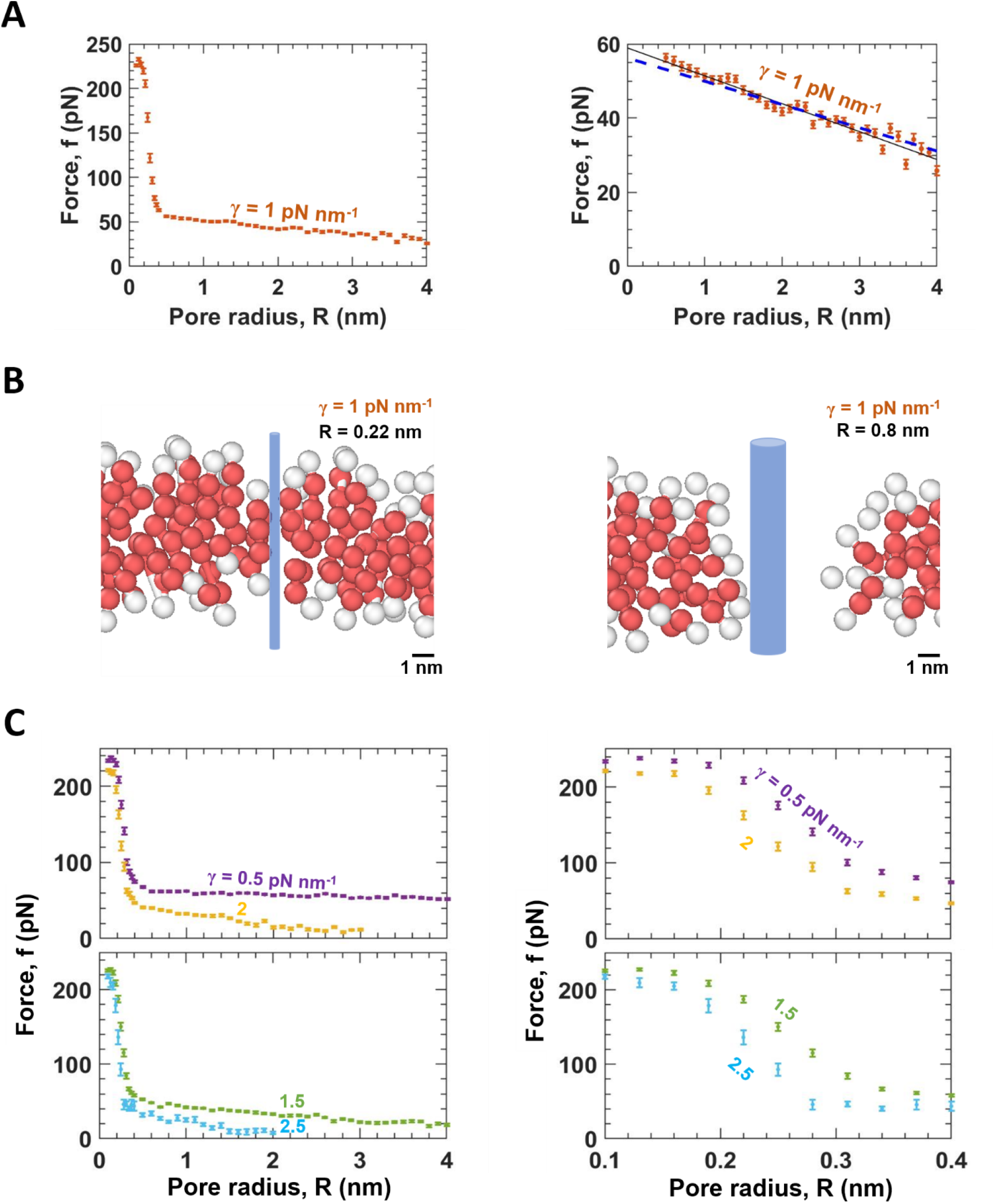
Force versus pore radius: small pores are hydrophobic and exert high force, large pores are hydrophilic and exert low force. **(A)** Time averaged force versus pore radius for a simulated bilayer at tension *γ* = 1 pN nm^−1^ (mean of *n* = 169 measurements per radius). For each pore radius, binned force (5 μs bins) was recorded in ~ 30 simulations each lasting ~5 – 50 μs for a total of *n* = 169 binned values, mean value. Right: Same data, showing only forces less than 60 pN. Best fit straight line for 0.6 nm < R < 4 nm (black line) has slope −7.53 ± 0.19 pN nm^−1^ (68% confidence interval, r^2^ = 0.95). Blue line has slope −2*πγ* = − 6.28 pN nm^−1^ predicted by classical pore model. Error bars: S.E.M. **(B)** Side view snapshots (vertical slices of thickness 1.7 nm) of pores with two radii from simulations in **(A)**. The small hydrophobic pore is lined mainly by hydrophobic tail groups. The large extended pore has a hydrophilic rim. **(C)** Mean force versus pore radius for four tensions (mean ± S.E.M., 181, 123, 94 and 32 measurements per radius from lowest to highest tension). Binning and simulation times as in **(A).** Right: same data, for radii 0.1 to 0.4 nm.

The small pores in the high force region were primarily lined with hydrophobic tail beads, Fig. 2B. Lipids at the pore edge tended to be “vertically” oriented (perpendicular to the bilayer plane) with their hydrophobic tails fairly tightly packed against the aqueous pore lumen enforced by the cylinder. By contrast, large pores in the low force region had curved rims formed by rotated lipids whose hydrophilic head groups lined the pore. Hydrophilic pores were much less tightly packed against the pore lumen and far more irregularly shaped, with gaps between the cylinder and the pore rim head groups (Figs. 1B, 2B). Relative to the cylinder enforcing the lumen, hydrophilic pores were much larger than hydrophobic pores.

Pore forces were lower at higher membrane tensions, Fig. 2C. The same qualitative behavior was observed over a range of tensions, 0.5 – 2.5 pN nm^−1^, with small hydrophobic pores exerting high forces at radii ~0.1-0.2 nm, but these forces were somewhat reduced at higher tension. Again, large hydrophilic pores exerted much smaller forces, but with higher tensions the force decreased more rapidly with pore size, with slope close to the −2πγ classical prediction (Fig. S3, Table S1).

### Pore line tension decreases with pore size, approaching a constant value for large pores

That small hydrophobic pores exert much larger forces than do large hydrophilic pores is easily understood from the work done to enlarge a hydrophobic pore, which proportionally increases the number of hydrophobic head groups in contact with the aqueous pore lumen. This is much more energetically costly than enlarging the hydrophilic rim of a hydrophilic pore.

This is most clearly seen in the pore line tension *τ*, defined to be *τ* = *∂F*/*∂S* for a pore of perimeter *S* = 2*πR* and area *A*. Since the free energy change for small changes is *dF* = *τ dS* − *γdA*, and given the definition of the pore force *f* ≡ *dF/dR*, it follows that the line tension is the force at zero tension divided by 2*π, τ* = [*f*/2*π*]_γ=0_. Fig. 3A shows the measured line tension *τ* versus pore radius *R*, which peaks at ~ 40 pN for small hydrophobic pores, then decreasing to a constant value *τ* ≈ 11 pN for pore radii above ~ 0.5 nm.

**Figure 3.**
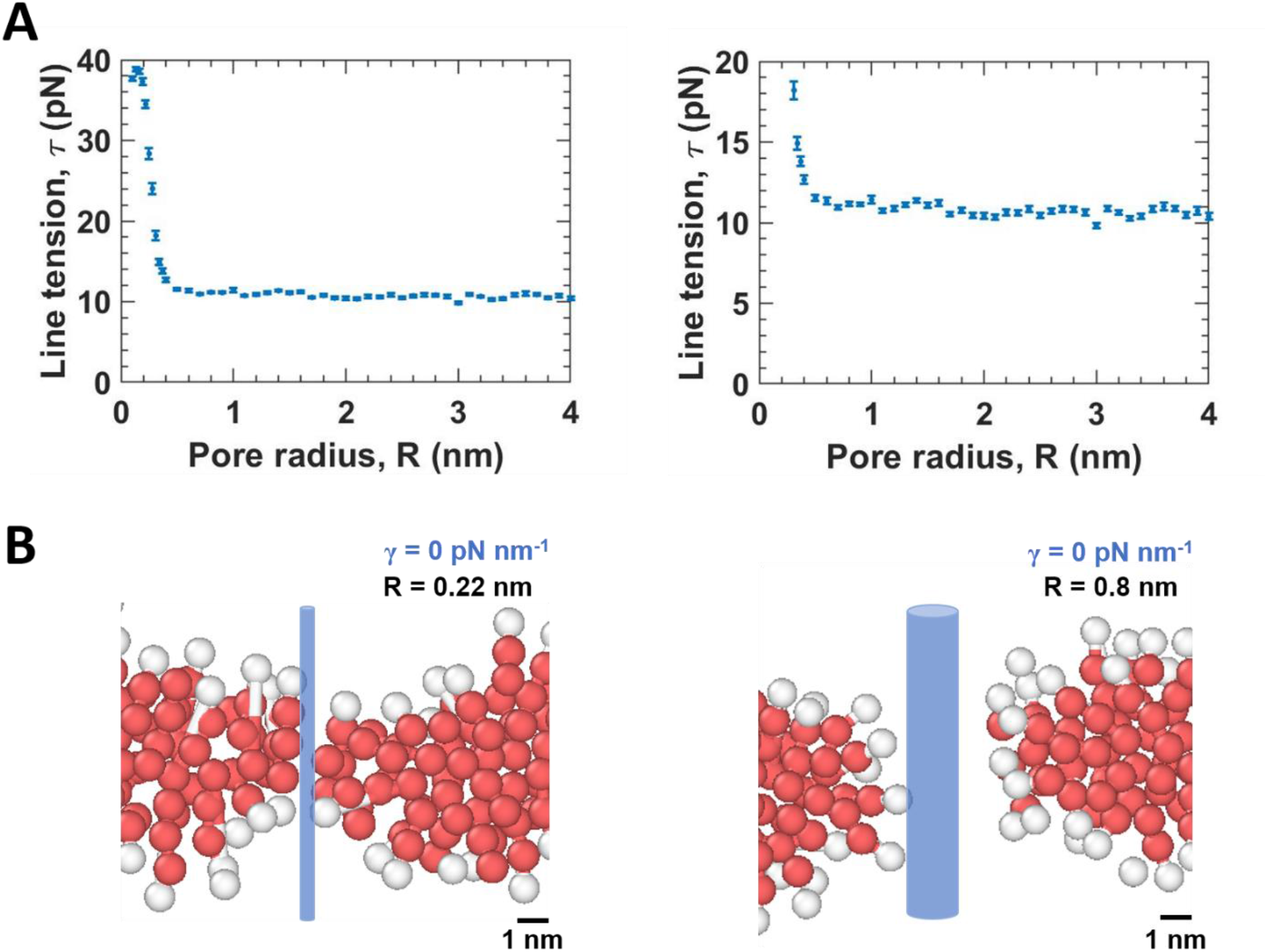
Pore line tension approaches a constant value for larger pores. **(A)** Pore line tension *τ* versus pore radius, calculated from the pore force for a tensionless bilayer divided by 2*π*. Mean of *n* = 228 binned force measurements per radius value, binned as for Fig. 2 Error bars: S.E.M. For larger radii, *τ* approaches a constant value 10.8 ± 0.3 pN (mean ± S.D.). Right: Same data, for *τ* < 20 pN. **(B)** Side view snapshots of pores with two radii from simulations in **(A)**, showing a small hydrophobic pore and a large hydrophilic pore.

Thus, pore line tension τ in general depends on pore size, but the concept of a constant *τ* becomes valid for larger pores: small pores have high *τ* due to their pore lumen-facing hydrophobic groups, while larger hydrophilic pores *R* > 0.5 nm approach a constant size-independent *τ* (Fig. 3B).

### Free energy vs pore size exhibits hydrophobic and hydrophilic pores but no metastable states

For each membrane tension, we integrated the force profile *f*(*R*) to obtain the pore free energy function, 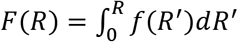. The free energy had a steep portion for small pores up to *R* ≈ 0.4 nm, and a shallower part for *R* > 0.4 nm, Figs. 4A-C. These two regions correspond to the high and low force regions described above which we associated with hydrophobic and hydrophilic pores, Figs. 2–3.

**Figure 4.**
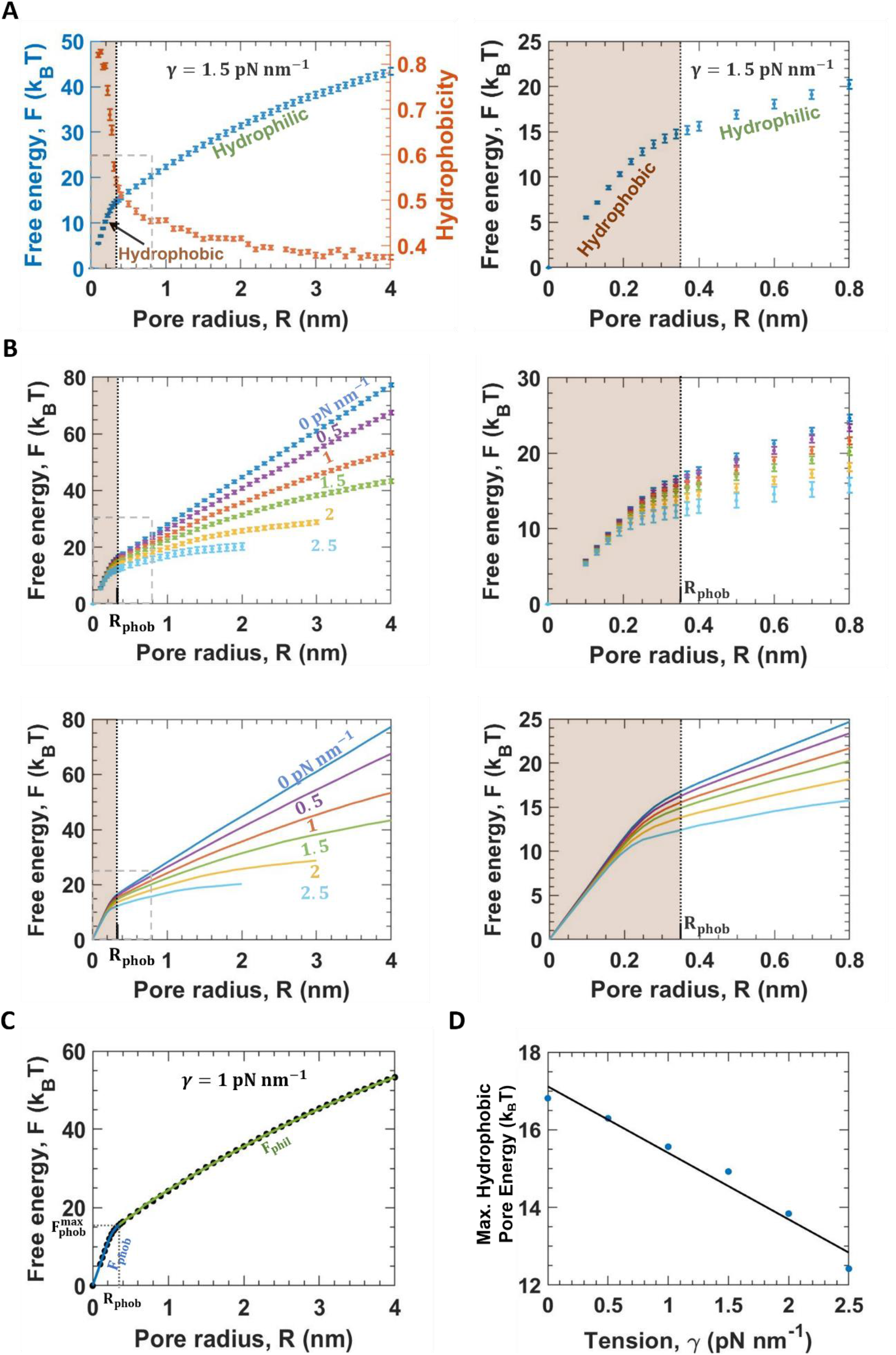
Free energy vs pore radius: a hydrophobic small pore domain, and a hydrophilic large pore domain. **(A)** Pore free energy versus pore size for simulated bilayers with tension 1.5 pN nm^−1^, obtained by integration of the corresponding force profile of Fig. 2C. Pore hydrophobicities are also plotted (see main text and Supplementary Information). The free energy curve has a steep region of high hydrophobicity pore (small pores), and a shallow region of low hydrophobicity or high hydrophilicity pores (large pores). The hydrophobic/hydrophilic boundary lies at *R*_phob_, the radius of the largest hydrophobic pore. Right: same free energy data, for pore radii up to 0.8 nm (dashed box at left). Error bars: S.E.M. Error propagation on the force profile S.E.M. values was used for free energy error bars. **(B)** Pore free energy versus pore size for the indicated membrane tensions. Right: same free energy data, for pore radii up to 0.8 nm (dashed box at left). Error bars: S.E.M. Error propagation on the force profile S.E.M. values was used for free energy error bars. The hydrophilic/hydrophobic boundary is shown at the average value of *R*_phob_ over all tensions (Table S2). Bottom row: same data as top row, but for clarity smooth curves through the mean values are plotted, without individual points. **(c)** Free energy fitting procedure illustrated for tension 1 pN nm^−1^. The hydrophilic portion is fit to *F*_phil_(*R*) (eq. (1) of main text), the classical pore model shifted by the hydrophilic rim radius, *δ*_rim_, with *δ*_rim_ and the constant *C* as fitting parameters. The largest hydrophobic pore radius *R*_phob_ is that where the hydrophobic free energy *F*_phob_(*R*) lies within *k*_B_*T* of the best fit *F*_phil_(*R*). **(d)** Maximum hydrophobic pore free energy 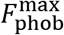 versus membrane tension from the fitting procedure of **(c)**. 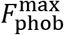 decreases approximately linearly with tension, with best fit slope 1/*γ*_phob_ where *γ*_phob_ = 0.58 ± 0.06, and intercept 17.12 ± 0.27 *k*_B_*T* (68% confidence interval, r^2^ = 0.96).

To quantify the pore type we measured pore hydrophobicity *H*, defined in terms of the lipid orientation angle *θ* relative to the Z axis, *H* = 〈|cos *θ*|〉 where the average is over lipids within 2 nm of the cylinder surface (see supplementary material, Fig. 4A). In the steep small pore region, *R* < 0.4 nm, hydrophobicity was high and approached a value ~ 0.8, while its value was much lower for larger pores and declined to *H*~0.4. This confirms that small pores are hydrophobic while large pores are hydrophilic.

Significantly, pore free energy profiles showed no evidence of long-lived metastable states (25,26) which would manifest as local free energy minima and regions of negative force. Our method cannot detect negative forces, but would register a metastable pore state by a finite interval of pore sizes with zero force and constant free energy. As there are no such regions in our simulations (Figs. 2–4) we definitively exclude metastability.

### Hydrophilic pores obey a classical free energy law shifted by the hydrophilic rim scale

Does the hydrophilic part of the free energy obey the classical model? Several findings conform to the classical picture: pore line tension *τ* approaches a constant, force curves have the classical slope, and free energy curves appear to approach a maximum for large pores, Figs. 2–4. However, it is immediately clear that the classical force prediction is not obeyed, *f* = 2*π*(*τ* − *Rγ*), since linear fits to simulated pore force profiles had tension-dependent intercepts extrapolated to zero pore size, inconsistent with the classical intercept of 2*πτ*, Fig. S3 and Table S2.

Thus we sought an improved description accounting for the approximately toroid-shaped hydrophilic rim seen in all large pores, Figs. 2–3. The extremely large rim thickness *δ*_rim_, of order 2nm, suggests that a hydrophilic pore with a minimum “waist” radius of *R* (set by the constraining cylinder) extends radially to a much greater effective distance *R* + *δ*_rim_. Then the energy released by membrane tension would be a greater advantage, but the line tension penalty a greater disadvantage, as described by the free energy

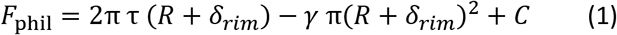

Using the measured value of pore line tension τ (Fig. 3), this form was a remarkably good fit to the measured free energy curves in the hydrophilic regions, with mean best fit rim thickness *δ*_*rim*_ = 2.02 ± 0.11 nm (mean ± s.d.), Figs. 4B, C and Table S2. The offset constant *C*, chosen to sew *F*_phil_ onto the hydrophobic part, *F*_phob_, decreased with tension (Fig. 4B, Table S2). We defined the end of the hydrophobic part as the smallest pore radius *R*_phob_ such that *F*_phob_ lies within *k*_B_*T* of *F*_phil_ (Fig. 4E).

In summary, the pore free energy is equal to *F*_phob_(*R*) in the hydrophobic region up to *R*_phob_, and *F*_phil_(*R*) of eq. (2) for larger pores. The maximal hydrophobic pore size *R*_phob_ = 0.35 ± 0.10 nm (mean ± s.d.) was almost independent of tension, while the maximal hydrophobic free energy 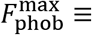 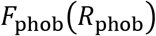 depended approximately linearly on tension

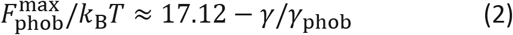

where *γ*_phob_ ≈ 0.58 pN nm^−1^.

### Membrane lysis times depend exponentially on membrane tension at high tensions

Membrane rupture is thought to occur by nucleation and growth of a membrane pore (12, 25). Having measured pores free energetics, we next directly measured rupture times versus membrane tension in simulations of 23 ×23 nm bilayers at tensions from 6 to 9.5 pN nm^−1^ (n=40 simulations per tension value). Following a rupture event the simulation was stopped, resulting in an aggregated time of ~1-18 ms per tension value.

The mean waiting time for rupture depended exponentially on tension, *t*_rup_~exp(−*γ*/*γ*_rup_) with *γ*_rup_ = 0.66 pN nm^−1^, decreasing from 3.7 ms to 23 μs over the tension range, Fig. 5A. Confirming the simple pore-mediated pathway, each rupture event was caused by nucleation of a pore that eventually grew without restraint, Movie S1.

**Figure 5.**
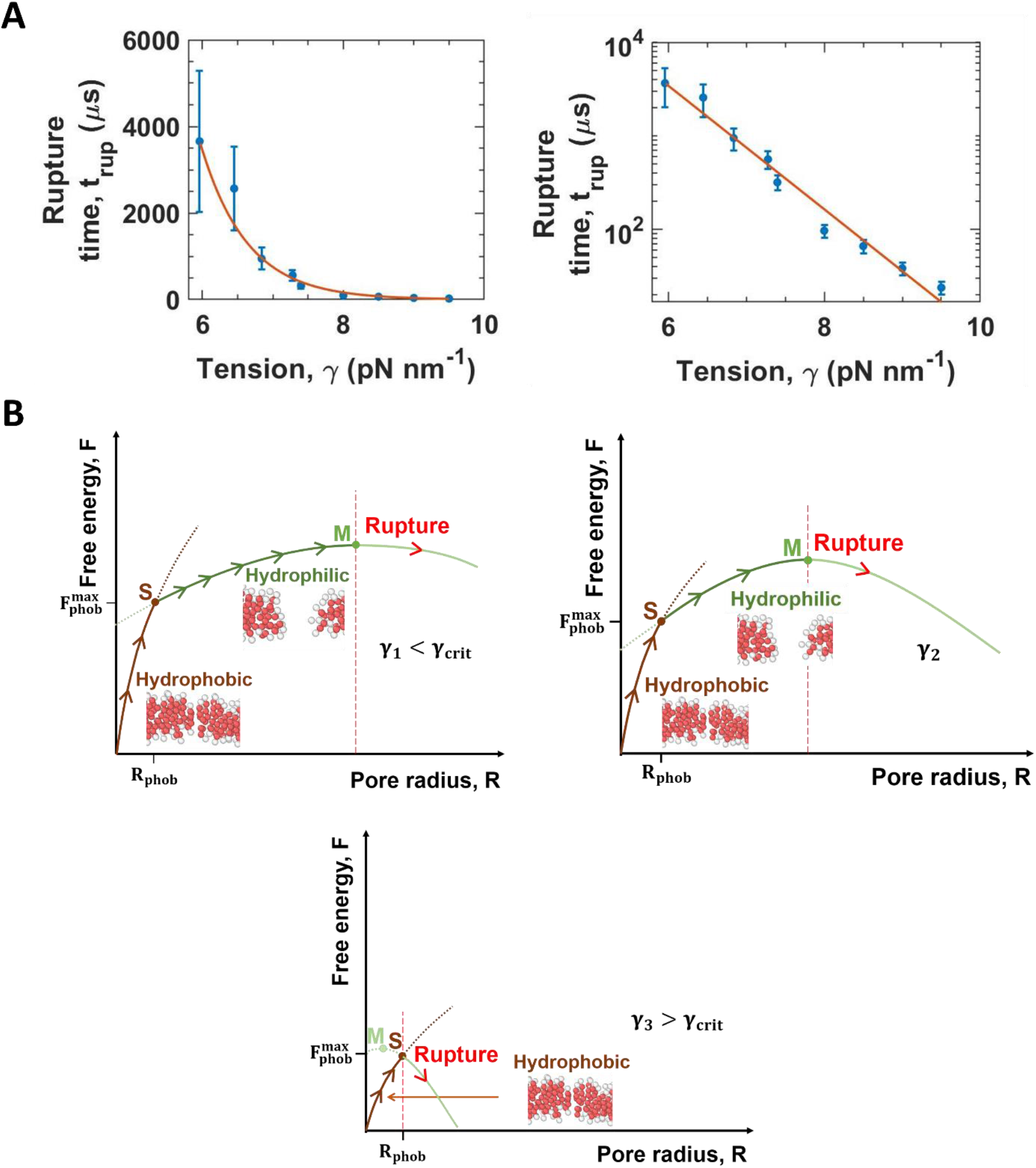
Membranes rupture via hydrophilic pores at low tensions but via hydrophobic pores at high tensions. **(A)** Mean rupture time versus membrane tension for simulated intact 23 × 23 nm bilayers without cylindrical constraint, shown in log-lin representation at right. Red curves: Best-fit exponential *t*_rup_ = *t*_0_ exp(−*γ*/*γ*_rup_), with *t*_0_ = 32 ± 26 s and *γ*_rup_ = 0.66 ± 0.05 pN nm^−1^ (errors are 68% C.I.). Error bars: S.E.M., mean values divided by 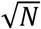, where *N* is the number of rupture events. **(B)** Schematic showing the pathway to membrane rupture emerging from the present study. Small pores in a membrane at subcritical tension *γ*_1_ lie on the hydrophobic free energy branch, *R* < *R*_phob_. Should a fluctuation grow a pore bigger than *R*_phob_ (~ 0.2-0.4 nm), it switches (point S) to the lower free energy hydrophilic branch. Growth to size *τ*/*γ*− *δ*_rim_ at the hydrophilic maximum (point M) gives rupture from a hydrophilic pore. Increasing tension (*γ*_2_ > *γ*_1_) moves M to smaller pore sizes. Above the critical tension *γ*_crit_ ≈ *τ*/*δ*_rim_ (~5 pN nm^−1^) M has crossed the hydrophobic branch (*γ*_3_ > *γ*_crit_) so that no pores large enough to accommodate the hydrophilic rim of size *δ*_rim_ are stable. Thus the membrane ruptures via a hydrophobic pore, from S to the unstable hydrophilic branch (negative slope).

Thus our simulations reproduce the exponential tension-dependence of rupture times observed experimentally by Evans et al. at high tensions (25). We note we were obliged to examine high tensions *γ* > 5 pN nm^−1^, as waiting times became unmanageably long for lower tensions.

### In the high tension exponential regime membranes rupture directly from a hydrophobic pore

Given the measured pore free energy profiles, Fig. 4, can we explain the exponential dependence of rupture times in simulations and experiments? Rupture entails pore growth to its limit of stability, the point of no return at the peak of the free energy profile *F*(*R*). Assuming simple mean field kinetics, the rupture time is governed by the free energy barrier, *t*_rup_~exp(*F*_max_/*k*_B_*T*) where *F*_max_ = πτ^2^/*γ* is the maximum of the hydrophilic free energy *F*_phil_ accurately describing our larger pore simulation data, eq. (1). Thus, the predicted tension dependence has the classical theory form, *t*_rup_~ exp(γc/γ).

This is far from the observed exponential behavior. The origin of this apparent discrepancy is that the hydrophilic branch does not exist at higher tensions. Indeed, the predicted maximally stable pore size at the hydrophilic free energy peak is *τ*/*γ* − *δ*_rim_, greatly reduced from the classical value by the pore rim thickness *δ*_rim_~ 2.0 nm, Table S2. Using *τ*~11 pN (Fig. 3A) it follows that this maximal hydrophilic pore size coincides with the edge of the hydrophobic pore region (*R*_phob_ ~ 0.35 nm) at a critical tension *γ*_crit_ = 4.7 pN nm^−1^, Fig. 5B. For higher tensions, all hydrophilic pores are unstable, and rupture occurs directly from a hydrophobic pore.

Thus, for high tensions the predicted rupture time is 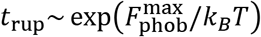. Using the linear tension dependence of the maximal hydrophobic free energy (eq. (2)), we predict a high tension exponential rupture regime,

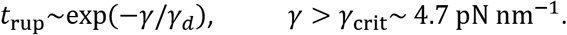

The predicted characteristic tension scale *γd*~0.58 pN nm^−1^ is close to that found in our rupture simulation runs, 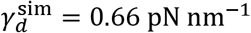.

## Discussion

Here, using highly coarse grained simulations to study pore energetics and dynamics, we found no evidence of macroscopically long-lived pore states in membrane bilayers. Free energies increased monotonically with pore size, with no indications of local minima, Fig. 4. Indeed, the presence of such anomalies is highly counterintuitive, given that typical lipid diffusivities ~ 1 μm^2^s^−1^ (35) naively suggest a nm sized pore would close in ~ μs timescales.

Metastable pore defects were proposed by Melikov et al. who detected ~ 1 s bursts of multiple ~ 1nm sized pore openings in BLMs subject to low electrical potential differences (26). The ~ 1s interval between bursts far exceeded the time between pore openings, suggesting an enhanced susceptibility to opening at some location persisted throughout a burst. They attributed this to electrically invisible metastable pre-pore defects with ~ 1s lifetimes of order the burst duration.

Similarly long-lived defects were invoked by Evans et al. to explain rupture data of ~20 μm GUV membranes subjected to tension ramp-up (25). Lower ramp up rate data was consistent with the classical rupture time prediction, *t*_rup_~ exp(*πτ*^2^/*γk*_B_*T*), but the high ramp up data implied an exponential dependence on tension for tensions *γ* > 5 – 10 pN nm^−1^. A modified classical pore model with a metastable ~ 0.6 nm prepore defect state was able to explain this regime: according to the model, the classical barrier to rupture is sufficiently reduced by high tensions that defect nucleation becomes rate-limiting. However, the model required the defects to have extraordinarily long lifetimes ~ 1s for their nucleation to be rate limiting, since even at these tensions the classical barrier of *πτ*^2^/*γ*~10 *k*_B_*T* remained high.

Our simulations used a well-tested highly coarse-grained force field lipid representation, shown to reproduce experimental bilayer bending moduli and phases (29–31). This was critical, as our simulated bilayer relaxation times were 1 – 5 μs, as seen from the ~ 1 μs pore memory time (Fig. S4A) and the ~ 5 μs tension memory time (Fig. S4B). Accordingly in most cases we pre-equilibrated bilayers for 20 μs; pores were clearly unequilibrated and the data unreliable on smaller timescales. Acceptable statistics required ~ 1 ms simulation times for a single free energy value (one pore size and membrane tension), Fig. 4, while rupture times were ~ 4 ms at tensions ~6 pN nm^−1^ (Fig. 5). Thus, meaningful equilibrium data is difficult to obtain from atomistic or MARTINI approaches, which were used to study pores over ~10-100 ns simulation timescales (28, 36). We note that other theoretical works have reported metastable pore states with local free energy minima of moderate 1-5 *k*_B_*T* depth from atomistic simulations (29–31), or depths ~ 10 *k*_B_*T* from a semi-analytical model (37).

Long ago it was proposed that electric fields drive electroporation of lipid bilayers by opening hydrophobic pores that mature into larger hydrophilic pores (12, 38). Here we showed that in equilibrium bilayers, small pores are hydrophobic and larger pores are hydrophilic with a large rim of width *δ*_rim_~ 2.0 nm. Above a critical tension the largest stable hydrophilic pore becomes smaller than the rim width, so stable hydrophilic pores no longer exist, Figs. 5B, C. Tension-driven membrane rupture then occurs directly from a hydrophobic pore with approximately exponential tension dependence of waiting times, Fig. 5A. Thus, without invoking macroscopically long-lived metastable pore defects, the observed high tension regime emerges naturally above a predicted tension ~ *τ*/*δ*_rim_~5 pN nm^−1^ similar to the experimental value 5 – 10 pN nm^−1^ (25).

## Acknowledgements

Research reported in this publication was supported by the National Institute of General Medical Sciences of the National Institutes of Health under award number R01GM117046 to B.O. The content is solely the responsibility of the authors and does not necessarily represent the official views of the National Institutes of Health. B. A. acknowledges support from NIH grant T32 GM008281. We acknowledge computing resources from Columbia University’s Shared Research Computing Facility project, which is supported by NIH Research Facility Improvement Grant 1G20RR030893-01, and associated funds from the New York State Empire State Development, Division of Science Technology and Innovation (NYSTAR) Contract C090171, both awarded April 15, 2010.

## Notes

### Competing Interest Statement

The authors have declared no competing interest.

