## Supplementary Information for "Membrane pore energetics and the pathways to membrane rupture"

### Supplementary methods

**Coarse-grained force fields for lipids developed in refs.** (1-3)

We use the force fields developed in refs. (1-3). Each lipid molecule is represented by 3 or 4 beads; we use the latter here. A lipid is a linear sequence of 1 head (h) bead followed by 3 tail (t) beads. Bonds exist between each successive pair of beads, and between the first and the fourth bead. Potentials of bonded and nonbonded interactions are given below and the parameters are in Table S1.

Non-bonded interactions

Repulsive potentials exist between all beads. This is of the form $V_{\mathrm{rep}}(r/b_{\mathrm{x}})$ where the subscript x can be 'hh', 'tt', or 'ht', $b_{\mathrm{x}}$ represents the length scale of repulsion, and $r$ is the distance between the centers of the two beads of interest.

$$V_{\mathrm{rep}}(x) = \begin{cases} 4\lambda(1/x^{12} - 1/x^6 + 1/4), & x \le 2^{1/6} \\ 0, & x \ge 2^{1/6} \end{cases}$$

$\lambda$ is an energy scale in the Cooke model. Based on this potential, a bead size can be defined as half of the smallest distance between centers of beads of the same type where they experience no repulsion. For head beads, this gives rise to a radius $r_{\mathrm{h}} = 2^{1/6} b_{\mathrm{hh}}/2$, and for tail beads the radius is $r_{\mathrm{t}} = 2^{1/6} b_{\mathrm{tt}}/2$.

Additionally, an attractive potential exists between tail beads of the form $V_{\mathrm{attr}}(r/b_{\mathrm{tt}})$

$$V_{\mathrm{attr}}(x) = \begin{cases} -\lambda, & x \le 2^{1/6} \\ -\lambda\ \cos\pi\left(x - 2^{1/6}\right)/2(w_c/\sigma), & 2^{1/6} \le x < 2^{1/6} + w_c/\sigma \\ 0, & x \ge 2^{1/6} + w_c/\sigma \end{cases}$$

$w_c$ represents the width of the attraction, and σ is the basic unit of length in the model.

Bonded interactions

FENE bonds are present between each successive pair of beads. The potential is given by

$$V_{\mathrm{bond}}(r) = -\frac{1}{2} k_{\mathrm{bond}} r_\infty^2 \log\left[1 - \left(\frac{r}{r_\infty}\right)^2\right],$$

where the bond stiffness $k_{\mathrm{bond}} = 30\lambda/\sigma^2$ and the divergence length $r_\infty$ is $1.5\sigma$.

A hookean spring ensures that the preferred distance between the centers of the first and the fourth bead is $6\sigma$, in order to restrict bending fluctuations. The potential is

$$V_{\mathrm{bend}}(r) = \frac{1}{2} k_{\mathrm{bend}} (r - 6\sigma)^2,$$

We used $k_{bend} = 10\lambda/\sigma^2$.

**Procedure used to open a pore in the initial bilayer**

Our initial condition is a crystalline bilayer. We then insert a cylinder with the following potential to open a pore for 2 μs. Then, we remove this cylinder and replace it with the cylindrical insert described in *Methods*, and then equilibrate for 20 μs.

$$V_{\text{init}}(r) = \begin{cases} 10\lambda, & r < R_\infty + r_{\text{h}} \\ h\big(r - (R_\infty + r_{\text{h}})\big), & R_\infty + r_{\text{h}} \leq r < R + r_{\text{h}} + 8\epsilon \\ 0, & r > R + r_{\text{h}} + 8\epsilon \end{cases}$$

where, $h(x) = 10\lambda \exp(-(x/0.62)^2)$.

**Procedure to measure hydrophobicity of the pore**

Hydrophobicity of the pore in Fig. 4A was calculated using snapshots taken every 2 µs from 6 simulations of 40 µs each for each radius (*n* = 120 snapshots per radius). Hydrophobicity was defined as the mean of the absolute value of the cosine of the angle made with the Z axis by the axis of all lipid molecules whose heads are within 2 nm of the surface of the constraining cylinder.

#### Supplementary figures

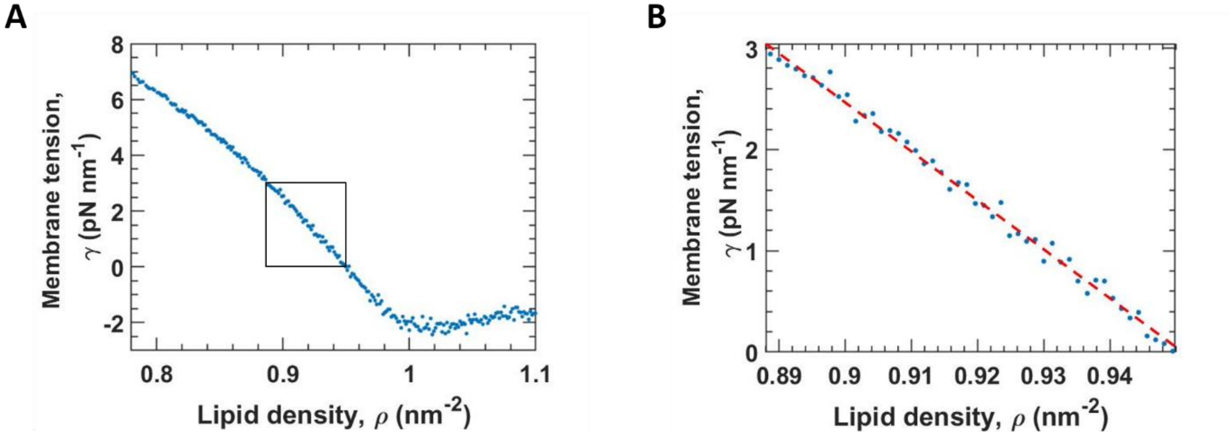


**Figure S1. Membrane tension in a simulated bilayer consisting of ultra-coarse-grained lipids decreases with lipid density for densities smaller than $\sim \mathbf{1\ nm^2}$. (A)** Membrane tension measured in our simulation versus lipid density. Each data point represents one simulation of a bilayer in the NVT ensemble in a cubic box of side 23 nm. Tensions were measured using the virial formula and averaged over the entire run. **(B)** Membrane tension versus lipid density data from the black rectangle in **(A)**. Dashed red line: best-fit straight line, with slope 48.3 ± 0.6 $\mathrm{pN\ nm}$, and y-intercept 46.0 ± 0.5 $\mathrm{pN\ nm^{-1}}$ (68% confidence interval, $r^2$ = 0.99).

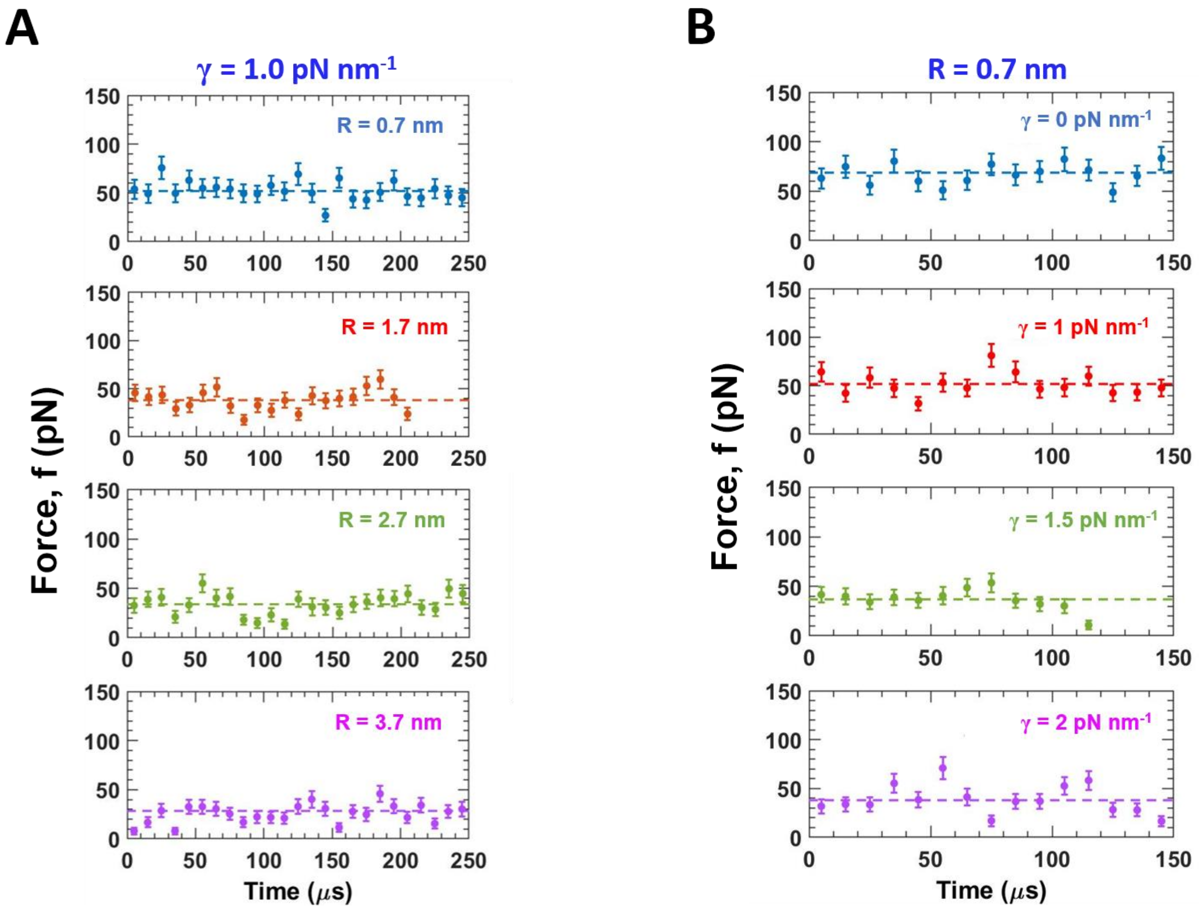


**Figure S2. Forces exerted by pores decrease with membrane tension and with pore radius for nanometer-sized pores (A)** Forces exerted by pores of given radii versus time in a membrane with tension $1\ \mathrm{pN\ nm^{-1}}$. Time-averaged forces (dashed lines) decrease with pore radius. **(B)** Forces exerted by pores of radius 0.7 nm versus time in membranes of given tensions. Time-averaged forces (dashed lines) decreases with tension. In both **(A)** and **(B)**, forces were measured every time step (0.2 ns) and binned over 10 μs (*n* = 50,000 measurements per bin), and each time course corresponds to one simulation. Error bars: S.E.M.

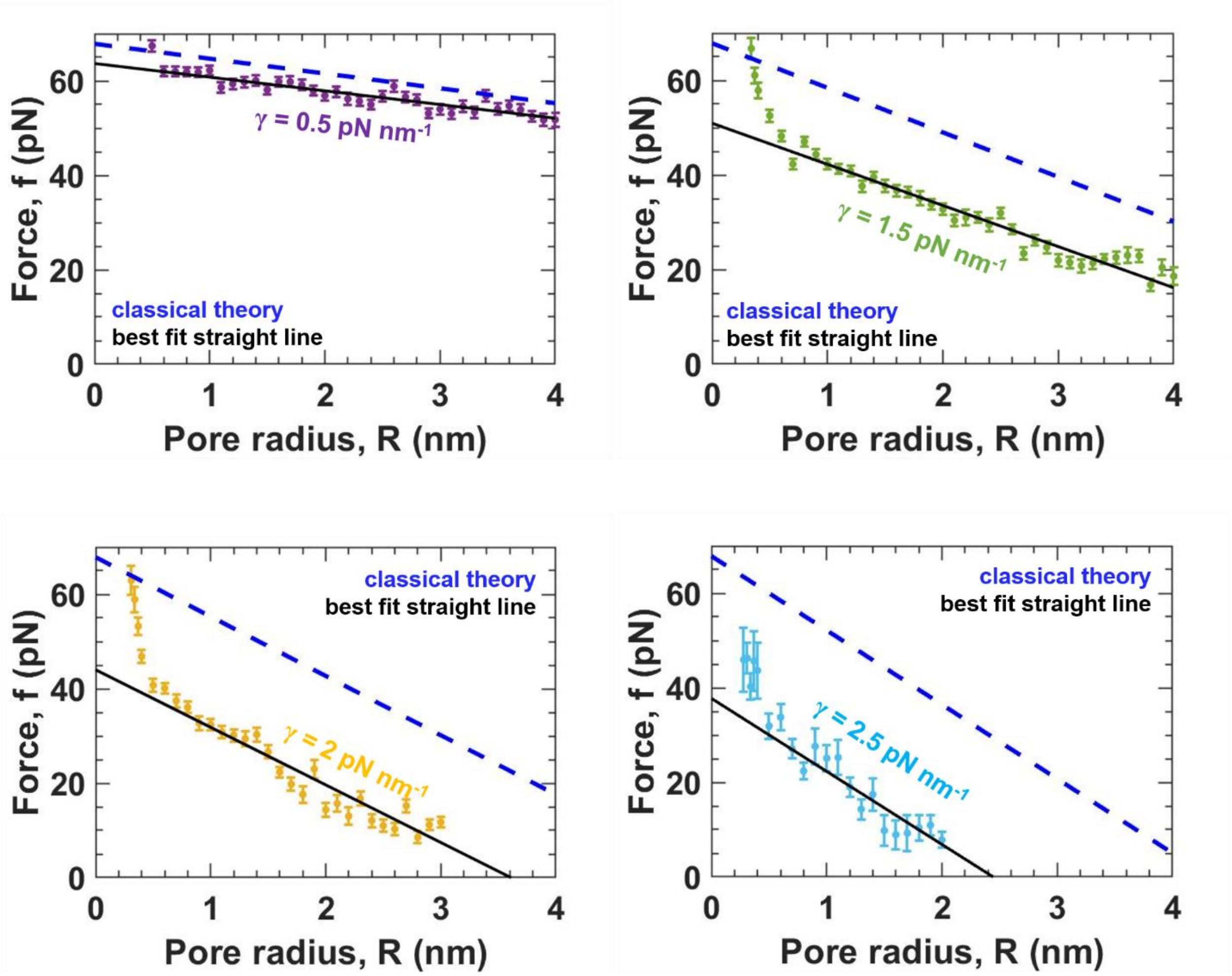


**Figure S3. Forces exerted by pores reduces with membrane tension.** Force versus pore radius with the given simulated tensions. Data is the same as that from Fig. 2C, but restricted to pore forces smaller than or equal to 70 pN. In this region, best-fit straight lines (black) have slopes consistent with the prediction of the classical model (blue dashed lines, predicted slope = $-2\pi\gamma$, Table S2). The y-intercept of the best-fit straight lines reduces with tension, unlike the prediction of the classical model that the y-intercept is equal to $2\pi\tau$ where $\tau$ is the line tension of the pore (Table S2). Best-fit straight lines were fit to radii larger than or equal to 0.6 nm. For the tension of 2 and 2.5 $\mathrm{pN\ nm^{-1}}$, the range of pore radii is comparatively smaller because all simulated bilayers ruptured at large pore radii. Error bars are S.E.M.

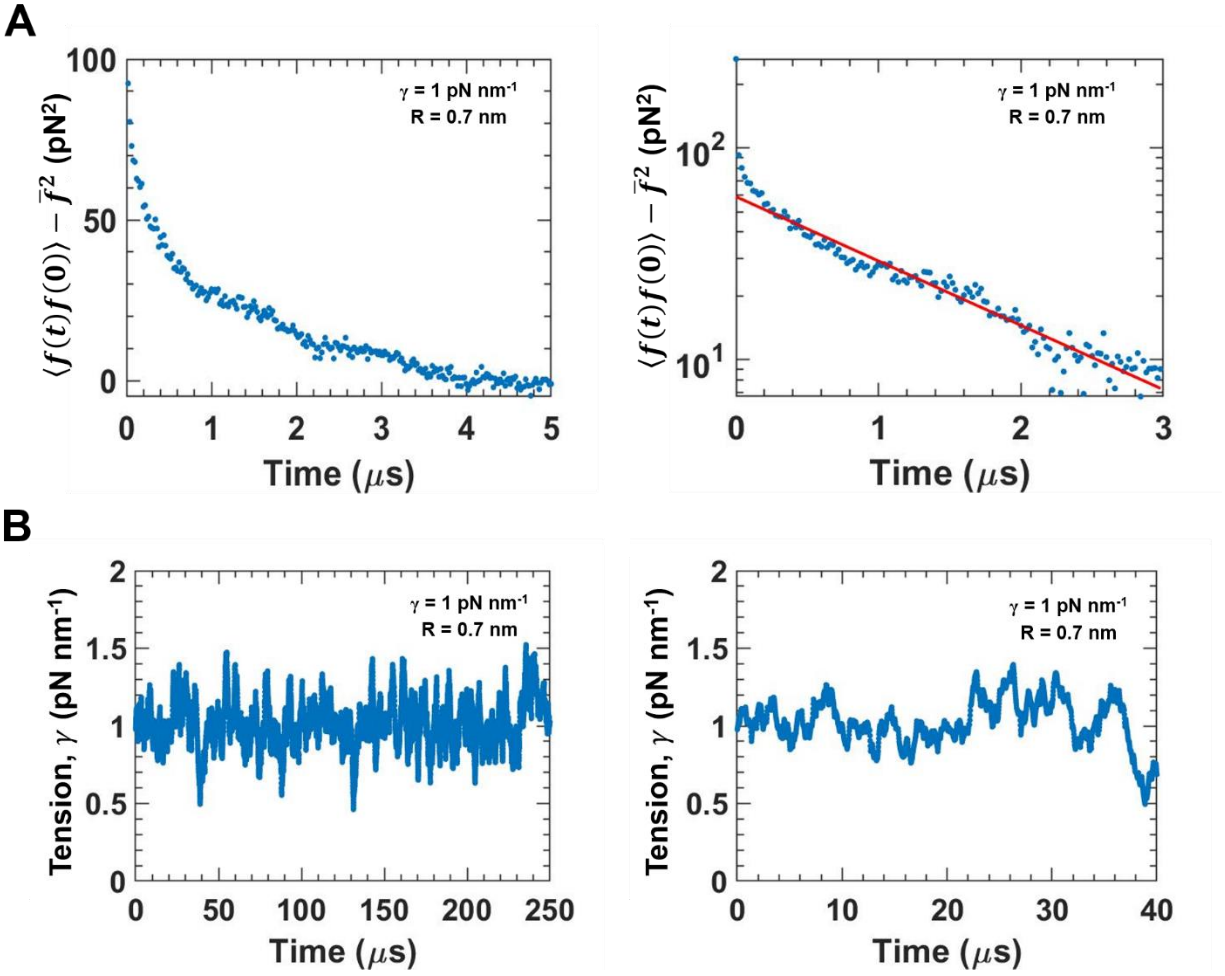


**Figure S4. Force exerted by pores and membrane tension measured from our simulations show microsecond correlation time scales. (A)** Temporal correlation function of the force exerted by the pore on the cylinder versus time for a simulated membrane with the given membrane tension and cylinder size (left), and the same data in a log-lin plot (right). Red line: Best-fit exponential function $\langle f(t)f(0)\rangle - \bar{f}^2 \sim exp(-t/\tau)$, with the correlation time $\tau = 1.42\ \pm 0.02\ \mu s$ (68% confidence interval). $\bar{f}$ refers to the time-averaged force. Forces were measured at every timestep (0.2 ns), and binned over a time interval of 20 ns. **(B)** Measured tension vs time over one entire simulation (left) and over a 40 μs period from the start of the simulation (right). Tensions were measured every 100 timesteps (20 ns) using the virial formula applied to patches of membrane of side 18 nm, one-fifth the side of the cubic simulation box (88 nm), and a moving average filter of size 2 μs was applied to the tension values. Tension shows a correlation time of $\sim 5\ \mu s$.

#### Supplementary Movie

**Movie S1.** Simulation of an intact bilayer with a tension $7\ \mathrm{pN\ nm^{-1}}$ and without any cylinder constraint, that eventually undergoes rupture. After a simulation time of ~35 $\mu s$, a pore gets nucleated and grows in an uncontrolled manner, leading to membrane rupture. The bilayer was simulated in a cubic box of side 88 nm, and a square patch of side ~40 nm is shown here.

| Symbol | Meaning | Value | Legend |
|---|---|---|---|
| $\sigma$ | Length scale in the Cooke model | 0.88 nm | (A) |
| $\epsilon$ | Width of cylinder repulsive potential $V_{\mathrm{cyl}}$ | 0.1 nm | (B) |
| $w_{\mathrm{c}}$ | Width of attractive potential between tail beads $V_{\mathrm{attr}}$ | 1.4 nm | (C) |
| $\lambda$ | Unit of energy in the Cooke model | 0.6 $\mathrm{k_B T}$ | (D) |
| $b_{\mathrm{hh}}, b_{\mathrm{ht}}$ | Head-head, head-tail repulsive length scale | 0.95 $\sigma$ | (D) |
| $b_{\mathrm{tt}}$ | Tail-tail repulsive length scale | $\sigma$ | (D) |

**Table S1. Parameters of the coarse-grained force field used to represent lipid molecules developed in** (1-3) **and parameters of the cylindrical insert.**

(A) Obtained by setting the measured bilayer thickness in the simulation to a typical experimentally measured value of 5 nm (4).
(B) Set as 1/8$^{\mathrm{th}}$ the diameter of a head bead
(C) Set equal to 1.6 $\sigma$ as in (1-3)
(D) Obtained from ref. (2).

| Measured tension $\gamma$ (pN nm$^{-1}$) | Best fit tension $\gamma_{\mathrm{fit}}$ (pN nm$^{-1}$) | Intercept/$2\pi$, β (pN) | Length scale $\delta_{\mathrm{rim}}$ (nm) | Hydrophobic radius $\mathrm{R_{phob}}$ (nm) |
|---|---|---|---|---|
| $0.09 \pm 0.17$ | $0.19 \pm 0.03$ | $11.18 \pm 0.07$ | n.a. | 0.29 |
| $0.54 \pm 0.16$ | $0.46 \pm 0.03$ | $10.14 \pm 0.08$ | 2.08 | 0.46 |
| $1.04 \pm 0.16$ | $1.19 \pm 0.03$ | $9.38 \pm 0.07$ | 1.80 | 0.31 |
| $1.51 \pm 0.16$ | $1.38 \pm 0.03$ | $8.11 \pm 0.08$ | 2.05 | 0.52 |
| $1.99 \pm 0.15$ | $1.94 \pm 0.06$ | $7.00 \pm 0.12$ | 2.09 | 0.24 |
| $2.46 \pm 0.15$ | $2.45 \pm 0.21$ | $6.00 \pm 0.29$ | 2.07 | 0.35 |

**Table S2. Best-fit parameters obtained from pore force and pore free energy versus pore radius in Figs. 2-4 and S3.** Best-fit tension values are obtained by dividing the slopes of the best-fit straight lines to force versus radius by $-2\pi$, as the prediction of the classical pore nucleation theory for the slopes of the force versus radius relation is $-2\pi\gamma$. $\delta_{\mathrm{rim}}$ was obtained using the relation $\delta_{\mathrm{rim}} = (\tau - \beta)/\gamma$ , where τ is the line tension (Fig. 3) and β is the y-intercept of the best-fit straight lines divided by 2π. The value of $\delta_{\mathrm{rim}}$ averaged over all tension values is $2.02 \pm 0.11$ nm (mean ± S.D.). The maximal hydrophobic pore size $\mathrm{R_{phob}}$ was obtained using the free energy of hydrophobic pores $F_{\mathrm{phob}}$ measured from our simulation, and the best-fit hydrophilic free energy $F_{\mathrm{phil}}$ (Figs. 4B-D). It was defined as the smallest pore radius such that $F_{\mathrm{phob}}$ lies within $k_{\mathrm{B}}T$ of $F_{\mathrm{phil}}$. The value averaged over all tensions is $R_{\mathrm{phob}}$ = $0.35 \pm 0.10$ nm (mean ± S.D.). The best-fit straight line to the free energy offset $C$ is $C/k_{\mathrm{B}}T = 20.9 - \gamma/(0.42\ \mathrm{pN\ nm^{-1}})$. Errors in the table are S.D. for $\gamma$, and 68% confidence interval for $\gamma_{\mathrm{fit}}$ and β. N.A. means not applicable.
